# Extracorporeal Photoimmunotherapy for Circulating Tumor Cells

**DOI:** 10.1101/013870

**Authors:** Gwangseong Kim, Angelo Gaitas

## Abstract

It is believed that metastasis through the cardiovascular system is primarily caused by circulating tumor cells (CTCs). We report an approach to eliminate circulating tumor cells from the blood stream by flowing the blood though an extracorporeal tube and applying photodynamic therapy (PDT). Chlorin e6 (Ce6), a photosensizer, was conjugated to antibody CD44 in order to target PC-3, a prostate cancer cell line. PC-3 cells were successfully stained by the Ce6-CD44 antibody conjugate. PDT was performed on whole blood spiked with stained PC-3 cells. As the blood circulated through a transparent medical tube, it was exposed to light of a specific wavelength generated by an LED array. A 2 minute exposure was sufficient to achieve selective cancer cell necrosis. Control studies showed no cell death.

## Introduction

Cancer metastasis is a major culprit for cancer death, given that it is responsible for over 90% of overall cancer deaths [1]. Metastasis occurs through the lymphatic or the cardiovascular system. In metastasis through the cardiovascular system, some primary cancer cells shed to the blood stream, circulate, and ultimately colonize other organs. Thus, circulating tumor cells (CTC) have a key role in cancer metastasis. A number of researchers have focused on detecting, enriching, and enumerating CTCs employing a number of mechanisms including: micro-fluidic separation devices [2-4], devices that rely on size exclusion by centrifugation [5, 6] or filtration [7, 8], immuno-magnetic separation [9, 10], and fluorescence activated cell sorting (FACS) technologies [2, 11] and several more techniques or combinations thereof. These technologies aim at diagnosis and are constrained by the small blood sample volume.

We hypothesize that removal of CTCs from the blood stream may reduce the chance of metastasis and the aggressiveness of existing tumors [12]. Recent studies report that there is indirect evidence that blood filtering, such as hemodialysis, might reduce cancer metastasis and the probability of cancer death by removing circulating tumor cells (CTCs) and other tumor growth factors from the bloodstream [13-15]. The rare frequency of CTCs and their presence in the circulatory system make traditional treatment modalities such as surgery, radiation, and chemo nearly impossible.

Extracorporeal filtration devices using leukocyte depletion filters have been used during tumor surgical procedures to remove tumor cells in order to reduce the risk of their dissemination [13-15], however these devices were not used to reduce metastasis post surgery. There have been efforts to remove or kill cancer cells using microtubes functionalized with antibodies, selectin, and cancer-specific tumor necrosis factor (TNF) - related apoptosis inducing ligand (TRAIL) with a capture and a kill rate between 30-41% [16, 17]. Recently a promising technique involved functionalizing circulating leukocytes with TRAIL and E-selectin adhesion receptor [18]. In a recent effort, our group functionalized a simple medical grade tube with EpCAM antibodies and successfully captured PC-3 cells in whole blood [19].

We propose an approach using extracorporeal photodynamic therapy (PDT, photoimmunotherapy in conjunction with antibody targeting). PDT requires three components, namely: oxygen, a photosensitizer, and light (mainly at the visible range). All these have to be present at the same time for the photosensitizer to be activated, generate reactive oxygen (mainly singlet oxygen, ^1^O_2_), and damage cells or tissues. Furthermore, the toxicity of the reactive oxygen species is localized at the cell in direct contact with it due to its short (< 100 nm) diffusion distance [20, 21]. These characteristics result in high specificity to target with near zero collateral damage to adjacent cells/tissues and result in an effective and safer treatment compared to conventional radiation and chemotherapy. In spite of these advantages, near-visible light can barely penetrate through tissue [22, 23], especially in the presence of blood (visible light absorber) and water (IR light absorber) and thus the application of PDT is mainly limited to diseases in opened/topical regions, including skin, head, neck, lung, teeth, etc.

We have developed a photosensitizer (Ce6) - antibody (anti-CD44) conjugate (Ce6- CD44 Ab conjugate) to selectively deliver the photosensitizing agent to CTCs (PC-3 cells in this case). An additional benefit to this technique is that the antibody can be safely cleared out of body by natural antibody degradation mechanisms within a few days [24]. PDT was performed by letting the blood flow through a thin transparent medical tube (Fig. 1). In this article, we demonstrate the proof-of-principle of this approach.

**Fig. 1.**
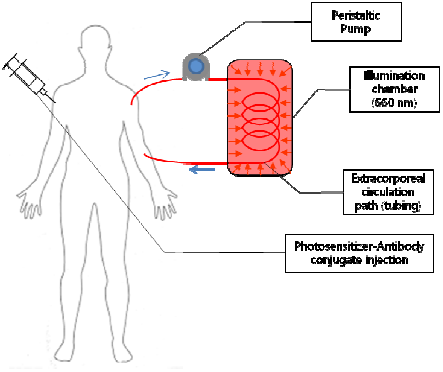
Schematic of the proposed device in operation. Photosensitizer-antibody conjugate is injected prior to PDT procedure. Blood circulation was guided by medical tubing with a peristaltic pump. Extracorporeal PDT is performed as the blood flows through the tube inside a reflective chamber. The treated blood is returned to body. All procedures can be in constant flow.

## Materials and Methods

### Conjugation of Ce6 to CD44 antibody

Chlorin E6 (Ce6) (Frontier Scientific) is a naturally occurring, commercially available photosensitizer that has excitation maxima in the far-red/near IR region (around 667 nm) and relatively high quantum efficiency. Because the Ce6 molecule has three carboxyl groups, it can be readily modified for chemical conjugation. Human CD44 antibody (BD Bioscience) was selected for the model human prostate cancer cell line, PC-3 (purchased from American Type Culture Collection (ATCC)). Expression of CD44 in PC-3 cell was previously reported [25] and confirmed experimentally by us. 2 mg of Ce6 was mixed with 6.5 mg of crosskinker, 1-Ethyl-3-[3-dimethylaminopropyl]carbodiimide hydrochloride (EDC) (Sigma-Aldrich) and 7.6 mg of sulfo-NHS (Pierce) in 1 mL PBS buffer at pH 7.4 (at 1:10:10 mole ratio respectively). The reaction ran for 2 hour at room temperature. Then, 50 μL of the product was added to 100 μL of FITC labeled human CD44 antibody solution. The conjugation reaction was run for 3 hours at room temperature with agitation. The reaction mixture was spin-filtered to remove unbound Ce6 residue at 4000 RCF for 100 min. The final product was resuspended in PBS, adjusting the total volume of 100 μL. The produced Ce6-CD44 Ab conjugate was stored at 4°C.

### Cell culture

The PC-3 cell lines were propagated in RPMI media supplemented with 10% fetal bovine serum (FBS) and 1% penicillin-streptomycin (PS). Passaging was done by trypsinization. Cell culture media, trypsin EDTA, and buffers were purchased from Life Technologies.

### Photodynamic therapy setup

PDT was performed using high power (maximum 100 W input) 660 nm LED array shown in Fig. 2. (a) (LEDwholesalers.com). Samples were placed within an aluminum foil covered styrofoam chamber shown in Fig. 2 (b) and Fig. 2 (c) and illuminated. The duration of illumination was studied to determine the optimal and minimum required time for PDT treatment.

**Fig. 2.**
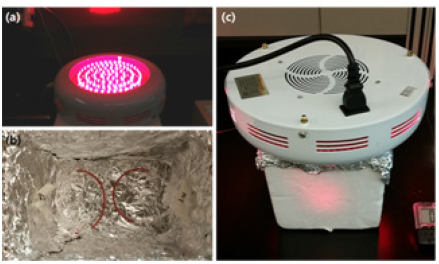
Extracorporeal PDT setup. (a) 660 nm LED array, (b) the aluminum foil covered chamber made from Styrofoam box with two tubes filled with blood. (c) PDT is performed by putting the LED array on top of the chamber. A 2 minute illumination was carried out to simulate extracorporeal blood circulation.

### PDT on PC-3 cells on a 12-well plate

PC-3 cells were cultured on a 12 well plate. Once the cell culture was confluent, 20 μL of Ce6-CD44 Ab conjugates were added and incubated for 1 hour. The cells were gently rinsed with warm DPBS three times to remove unbound excess Ce6-CD44 Ab conjugates and the plate was filled with 1 mL RPMI medium. The staining of the Ce6-CD44 Ab conjugates was confirmed by the fluorescence of FITC label on CD44 antibody. 10 μL Calcein AM (Life Technologies) was added to the cells and incubated for 20 min. The extra Calcein AM was removed by rinsing three times and the plate was refilled with 0.5 mL warmed whole blood (Innovative Research). PDT was performed by putting the 12-well plate in the PDT chamber and illuminated for 2 min. The weakening of the Calcein AM florescent signal is an indication of cell death. We monitored the florescent signal every 15 minutes for 2 hours. Three control experiments were performed in parallel. The first control experiment involved PC-3 cells stained by Ce6-CD44 Ab conjugate in culture media and illuminated by the LED (positive control). The second control included PC-3 cells stained by Ce6-CD44 Ab conjugate but not illuminated by the LED (no light - negative control). The third control consisted of PC-3 cells that were illuminate by the LED but were not stained by Ce6-CD44 Ab conjugate (no conjugate - negative control). The outcome of PDT was monitored by fluorescence imaging with an Olympus IMT-2 inverted microscope connected to a Zenoptik MF Progres camera using Progres Capture Pro software.

### PDT on PC-3 cells in a tube

PC-3 cells were suspended using 0.25% trypsin EDTA. The number density of floating cells was estimated by hemocytometry. 300,000 cell / mL were used for the study. 20 μL of Ce6-CD44 Ab conjugates were added to 1 mL cell suspension and incubated for 1 hour. The cells were pelleted by centrifugation at 1000 RPM for 5 min. The supernatant was removed and the cells were resuspended in 1 mL of fresh RPMI medium. The staining by Ce6-CD44 Ab conjugates was confirmed by fluorescence microscopy based on the FITC label present on the CD44 antibody. The PC-3 cells were stained by Calcein AM for 20 min to monitor cell death. The cell suspension was centrifuged at 1000 RPM for 5 min and the supernatant was removed. The cell pellet was resuspended in RPMI medium to a final density of about 50,000 cells / 20 μL. 20 μL of these cells were spiked into 100 μL warmed whole blood. After mixing, a 15 cm Silastic polydimethylsiloxane (PDMS, Silicone) tubing (Fisher Scientific) was filled with the PC-3-blood mixture and illuminated for 2 min. The blood specimens were collected. The fluorescence of Calcein AM labeled PC-3 cells was monitored at 0 min (before illumination), 1 hour, and 2 hours respectively using hemocytometry on a fluorescence microscope.

Two control experiment were performed. The first control included illuminated PC-3 cells without Ce6-CD44 Ab conjugate (no conjugate - negative control). The second control included PC-3 cells stained with Ce6-CD44 Ab conjugate but with exposure to light (no light - negative control). The reduction in the strongly fluorescing cells was analyzed to determine the efficacy of PDT.

## Results

### PDT on PC-3 cells on a 12-well plate

PC-3 cells were grown confluent on the substrate of a 12-well plate. After 1 hour of incubation with the Ce6-CD44 Ab conjugate, positive staining was confirmed by monitoring the FITC label on the CD44 antibody (Supplementary Fig. 1). The cells were stained with a cell viability indicator, Calcein AM. Illumination with the LED was performed in the presence of either 0.5 mL of whole blood or, in the case of the positive control, in 0.5 mL RPMI media. The results are summarized in Fig. 3. In the positive control in RPMI media nearly all of the PC-3 cells were killed within the first 15 minutes (in the 15 minutes we are including the 2 minutes of illumination) without any noticeable survival. This result demonstrates that the Ce6-CD44 Ab conjugate is a highly potent photosensitizer reagent with target specificity. In the presence of whole blood cell death was slowed down and a small population of cells survived after two hours. The most probable reason for these results is that hemoglobin in blood is blocking some of the light from reaching through to the cells at the bottom of the plate. Both negative controls showed negligible reduction in Calcein AM staining and thus there was no cell death observed.

**Fig. 3.**
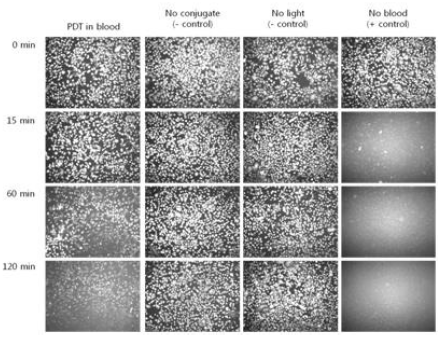
Results of photodynamic therapy after 2 minutes illumination on a plate. Cancer cells are stained with calcein AM. The left hand side “PDT in blood” column shows a slowdown in cell death compared to the “no blood” column because light is absorbed by blood. The negative controls do not exhibit cell death.

### PDT on PC-3 cells in a tube

PC-3 cells spiked in whole blood were inserted in a thin laboratory tube to evaluate the efficacy of extracorporeal therapy. About 50,000 PC-3 cells pre-stained with Ce6-CD44 Ab conjugates and Calcein AM were spiked in 100 μL of whole blood. The cell killing effect was studied by monitoring the Calcein AM fluorescence in the presence of blood and also quantified by performing fluorescence imaging on a hemocytometer. The sample that contained the Ce6-CD44 Ab conjugate, and was illuminated, demonstrated successful cell-death. The samples showed similar initial density of fluorescing PC-3 cells prior to illumination (Fig. 4). We monitored cell death after 60 min and 120 min and the results showed significant reduction in fluorescing cells for the cells that were treated with the Ce6-CD44 Ab conjugate and illuminated (Fig. 5). The two negative controls (one with conjugate but without illumination, and the other without conjugate but with illumination) did not exhibited noticeable change in the density of fluorescing cells within the 2 hour of observation (Fig. 5). PDT in a tube appears to be more effective in killing cells than in the 12-well plate.

**Fig. 4.**
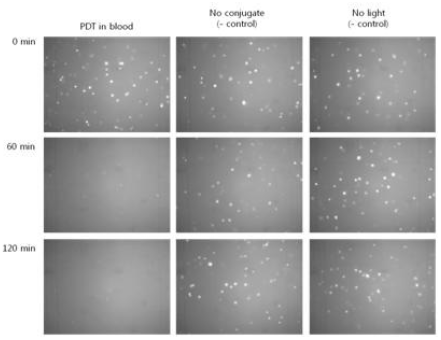
Results of photodynamic therapy in a tube with 2 min illumination. The tube’s diameter is 1.02 mm, which is within the penetration depth of light given that the tube is illuminated from all directions, for this reason there is faster cell death compared to the results in media, as shown in the left hand side “PDT in blood” column.

**Fig. 5.**
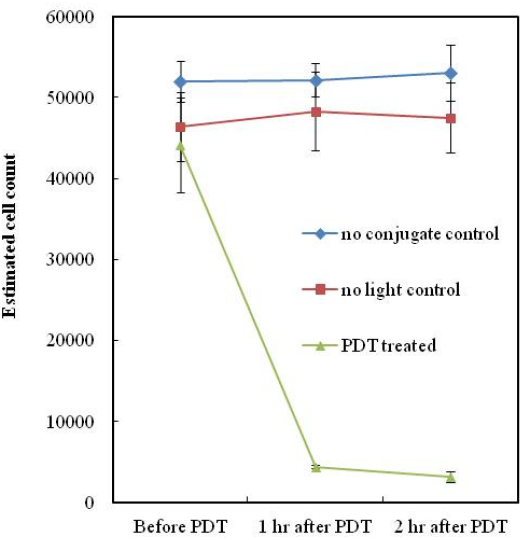
The quantitative analysis of PDT outcome for PC-3 cells in tube. (n=3, data represent mean ± standard error)

## Discussion

PDT is an effective alternative treatment modality which addresses several of the drawbacks of conventional treatments in cancer and in other diseases. However, the absorption of visible light by blood (especially due to the red blood cells’ hemoglobins) significantly reduces the penetration of light through tissue. This is evident in our results shown in "PDT in blood" column of Fig. 3, where PC-3 cells were cultured on a 12-well plate. Following a 2 minute illumination 2 hours later the some cells were still alive. In contrast, the cells in media were completely wiped out by the 2 minute illumination in the absence of blood within the first 15 minutes. This result clearly showed how much blood can hamper the PDT’s efficacy.

The PDT results, from Fig. 4, where a tube was used exhibited improved efficacy. We believe that the improvement came from the utilization of the tube. The tube used in this study is a transparent PDMS (silicone) tube with 1.02 mm inner diameter. Since the light came from all the directions surrounding the tube in the reflective chamber, the thin diameter of tube allowed for nearly the entire sample to be within the penetration depth of light. Allowing for more exposure to light resulted in better outcomes with PDT.

The experimental parameters used in this investigation, such as the choice of photosensitizer, the illumination time, the antibody, the type of tube material, size of tube (length and diameter), the light source etc. need fine tuning. These parameters should be further studied and optimized to obtain maximum efficacy of PDT, especially with consideration to in vivo constant flow conditions.

A similar approach was suggested previously by Edelson et al. [26] where the "extracorporeal photopheresis (ECP)" concept was first reported. However, in this technology the blood is collected and separated by apheresis and buffy coat was treated with UV light and reinjected to the body. This technology is used to treat cutaneous T-cell lymphoma and graft-versus-host diseases (GVHD) in organ transplantation and is currently approved by the FDA. Our approach has the benefits of not requiring blood separation processing and of utilizing light in the far-red/near infrared wavelength that has deeper tissue penetration depth. In addition, the photosensitizer-antibody conjugates can be used as an imaging contrast to detect metastasized cancers, allowing other treatment modalities, including endoscopic photodynamic therapy. What is more it would be interesting to examine the possibility of targeting the lymphatic system. Finally, this concept can be translated to target other diseases.

---

**Supplementary Fig. 1.**
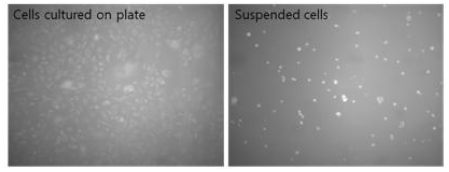
Evidence of positive staining PC-3 cells by Ce6-CD44 Ab conjugate in 12 well plate (left) and suspension (right).

